# The genome of the *Delisea pulchra*: a resource for the study of chemical host-microbe interactions in red algae

**DOI:** 10.64898/2026.03.31.715562

**Authors:** Simon M. Dittami, Jennifer Hudson, Loraine Brillet-Guéguen, Elizabeth Ficko-Blean, Gwenn Tanguy, Sylvie Rousvoal, Erwan Legeay, Gabriel V. Markov, Ludovic Delage, Olivier Godfroy, Erwan Corre, Jonas Collén, Catherine Leblanc, Suhelen Egan

## Abstract

**Background:** Red macroalgae (Rhodophyta) are ecologically and economically important marine primary producers, yet genomic resources for most species remain scarce. *Delisea pulchra*, a temperate red alga known for its halogenated furanone-based chemical defenses, serves as a model for studying algal-microbe interactions, antifouling mechanisms, and disease dynamics.

**Results:** Here we present a high-quality genome assembly of this species. The nuclear genome comprises 134 Mbp across 271 contigs with an N50 of 1.47 Mbp and encodes 13,387 predicted protein-coding genes. Comparative genomics with other red algae revealed expansions in gene families involved in DNA methylation, and oxidative stress responses, including glutathione S-transferases and superoxide dismutases. Analysis of glycosyltransferases, sulfatases, and sulfurylases implicated in galactan biosynthesis suggests *D. pulchra* possesses a complex and potentially novel extracellular matrix. We also identified several vanadium haloperoxidases (vHPOs), heme-dependent haloperoxidases (hHPOs), and two type III polyketide synthase (PKS) genes unique to the *D. pulchra*, which together represent promising candidate genes for bromofuranone production.

**Conclusion:** The *D. pulchra* genome provides a foundation for molecular investigations into defense, signaling, and host-microbe interactions. It has been deposited at the European Nucleotide Archive under accession number PRJEB101077. All datasets, annotations, and interactive tools for exploring the genome are also available through the Rhodoexplorer portal at https://rhodoexplorer.sb-roscoff.fr.

## Introduction

Macroalgae (seaweeds) are a diverse group of photosynthetic organisms that are critical to the ecological functioning of temperate marine habitats. As primary producers, macroalgae provide food and habitat for macroscopic and microbial life forms and provide a foundation supporting broader trophic interactions in the ocean. However, macroalgal communities globally are currently under considerable threat from the effects of anthropogenic activity, such as climate change, which has been linked to an increase in disease outbreaks as well as localised extinctions (Hanley et al. 2024).

Rhodophyta (red algae) represent a large and diverse monophyletic lineage of algae, comprising over 7,000 species of micro- and macroalgae (Guiry, M.D. & Guiry 2025). In addition to their ecological roles, species of Rhodophyta are highly regarded for their economic value as a food source (e.g. nori). They are also a rich source of complex polysaccharides (such as agars and carrageenans), which have applications across the food, biotechnology, cosmetic, and pharmaceutical industries. Red algae additionally synthesise and secrete a broad range of unique secondary metabolites such as terpenoids, phenolics, and flavonoids. Ecologically, these compounds function to deter disease-causing microorganisms, as well as herbivores and fouling organisms, while also serving as a reservoir of bioactive compounds with potential applications in novel drug discovery and sustainable agricultural practices (Carpena et al. 2023).

The red macroalga, *Delisea pulchra* (Bonnemaisoniales) (Fig. 1), is found along temperate coastlines of Australia, New Zealand, Japan, and sub-Antarctic regions. This seaweed has been studied extensively over the last two decades due to its unique defensive chemistry (Harder et al. 2012). *Delisea pulchra* produces halogenated furanone molecules that not only have anti-fouling and anti-herbivory activities (Dworjanyn et al. 2006), but are renowned as natural antagonists of bacterial cell-cell communication signalling (Rasmussen et al. 2000). More recently, *D. pulchra* has been used as a model to elucidate the molecular basis of microbial diseases in red algae (Zozaya-Valdés et al. 2017), characterise algal pathogen response mechanisms (Hudson et al. 2022) and develop disease prevention strategies such as the implementation of probiotic microorganisms (Li et al. 2022).

**Figure 1.**
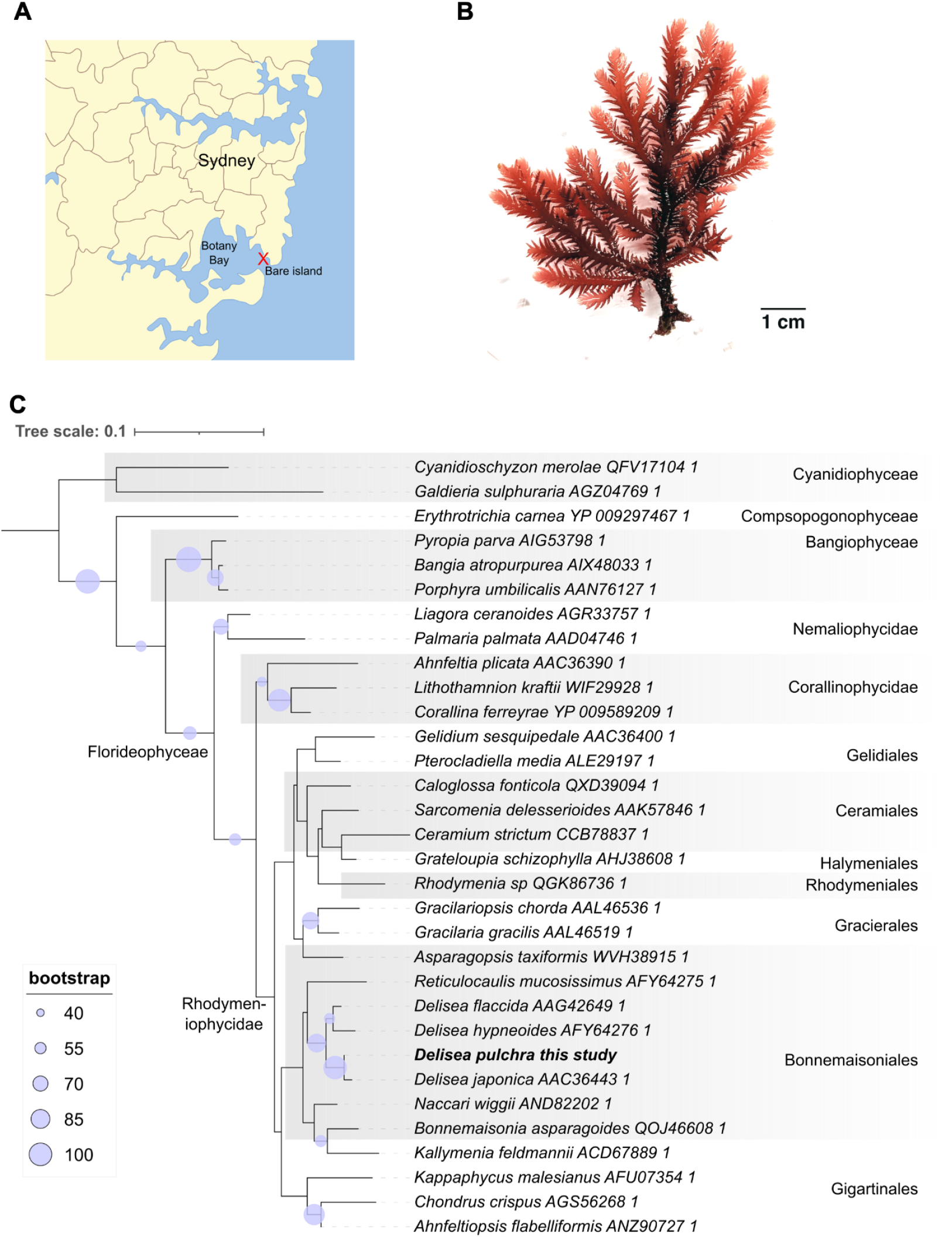
A) Sampling location of *Delisea pulchra* individuals (map by New South Wales local government under the CC-by-sa 3.0 license) B) Photo of a juvenile *D. pulchra* individual under laboratory conditions C) Maximum likelihood phylogenetic tree of selected red algae based on the rbcL gene. The tree was generated on ngphylogeny.fr (Lemoine et al. 2019) using MAFFT for the alignment, BMGE for curation (461 conserved positions), Phyml for tree reconstruction (LG model, 100 bootstrap replicates), and iToL for tree visualisation. The Cyanidiophyceae were considered the outgroup and used to root the tree.

Despite their global significance, red algal genomes remain under-characterised, limiting our capacity to understand and enhance disease resistance and environmental resilience in algal communities. This also presents a bottleneck to exploiting this genetic reservoir for industrial applications. Here, we present a genome assembly of the model red alga *D. pulchra* to explore the genetic basis of its unique metabolic capacities and to further our knowledge of host-microbe interactions in red algae.

## Materials and Methods

### Biological material, DNA extraction, and sequencing

Two individuals, likely sporophytes and approximately 30 cm in height, were collected from Bare Island, Sydney, Australia (-33.992030, 151.232149) at a depth of 10 meters in August 2020 and July 2022. The samples were immediately returned to the laboratory, where they were rinsed in sterile seawater followed by sonication for 10 minutes to remove visible epiphytes. Samples were stored in TNES urea buffer (10 mM Tris-HCl pH 7.4, 120 mM NaCl, 10 mM EDTA pH 8.0, 0.5% SDS, 4 M urea, pH = 8) until further use.

The first sample was extracted using a protocol based on the Macherey-Nagel Nucleobond® AXG columns. Approximately 200 mg of algal tissue were ground in liquid nitrogen, then incubated in 550 µL of Lysis Buffer CF and 10 µL Proteinase K (20 mg/mL) at 65°C for 1h, adding 10 µL of RNase (25 mg/mL) after 30min. The lysate was centrifuged at 4000g for 30 min, and the supernatant was then purified on AXG500 columns according to the manufacturer’s instructions, starting at step 2 of the Isolation of Genomic DNA protocol. The final pellet was resuspended in 300 µL of the elution buffer and stored at -80°C until library preparation using the Illumina DNA Prep workflow (ref. 20060060). Sequencing of sample one was carried out at the Genomer platform of the Station Biologique de Roscoff (France) on an Illumina MiSeq using a V3 MiSeq Reagent Kit and 2x75 bp read length.

For the second sample, 1 g of preserved tissue was frozen in liquid nitrogen, then ground finely in a mortar with a pinch of sterile sand. The extraction protocol was adapted from Fauchery et al. (2018). Briefly, ground tissue was resuspended in 17.5 ml of CTAB based lysis buffer (0.1 M Tris-HCl, 0.75 M NaCl, 0.75% CTAB, 20 mM EDTA, 0.13 M Sorbitol, 0.75% Sarkosyl, 0.01% PVP, 2.5mg Proteinase K), then incubated for 30 min at 65°C. Cell debris were precipitated on ice for 30 min by adding 5.75 ml of potassium acetate (5M). The supernatant was collected after centrifugation at 5000g for 20 min at 4°C. After extraction with 1 volume of chloroform:isoamyl alcohol (24:1), the polysaccharides were precipitated by adding 0.3 volumes of ethanol, then extracted with 1 volume of chloroform:isoamyl alcohol (24:1). Finally, genomic DNA was precipitated with 1 volume of isopropanol, desalted, dried, then eluted in 200ul of TE buffer (10mM Tris, 1mM EDTA) pH8.0. It was purified on a Qiagen Genomic-tip 500/G column, according to the manufacturer’s instructions. Lastly, the genomic DNA was eluted with 15 ml of prewarmed (50°C) QF buffer, then precipitated. Library preparation was carried out using the SMRTbell Prep kit 3.0 (ref. 102-182-700), and PacBio sequencing was performed on the SMRTCell Sequel II sequencer at the Gentyane platform at Clermont-Ferrand (France).

### Assembly and removal of non-target sequences

Adapter sequences remaining in the PacBio CSS reads were removed using Cutadapt version 2.8 (Martin 2011). The cleaned reads were used to perform de novo genome assembly using HiFiAsm version 0.15.1 and default parameters (Cheng et al. 2024). Plastid and mitochondrial genomes were assembled with NOVOPlasty version 4.3.4 using the MiSeq reads and the *Chondrus crispus* plastid (HF562234.1) and mitochondrion (NC_001677.2) as references (Dierckxsens et al. 2020).

Removal of non-target sequences was performed using the Anvi’o pipeline version 8 (Eren et al. 2020). Briefly, the MiSeq reads were mapped to the assembly using Bowtie version 2.5.1 (Langmead et al. 2009), while PacBio CSS reads were mapped to the assembly using Minimap 2.24 (Li 2018). Contigs and profile databases were generated with Anvi’o and COG and HMM domain information was added using the “anvi-run-ncbi-cogs” and “anvi-run-hmms” functions, respectively. Taxonomic information was predicted for all gene calls using Kaiju version 1.9.2 and the Kaiju nr_euk database from May 10th, 2023. Finally, binning was carried out manually. The bin corresponding to the *D. pulchra* genome was separated from the other bins since it was the only bin that had substantial coverage by MiSeq (sample 1) and PacBio reads (sample 2).

### RNA-seq mapping and structural annotation

Deduplified RNAseq reads were obtained from (Hudson et al. 2022) and mapped against the clean *D. pulchra* genome using Tophat version 2.1.1 (Kim et al. 2013). All but four genomic contigs had mapping RNAseq reads, and these four contigs were compared to the NCBI nt database (4/01/2024) using the dc-megablast algorithm, to manually confirm to confirm these were not belonging to microbial contaminants (e>0.05). RepeatModeler 2.0.3 and RepeatMasker version 4.1.2 (Flynn et al. 2020) were used to generate a database of repeated elements and soft-mask these before predicting coding sequences. Gene prediction was then carried out using Braker 3 (Gabriel et al. 2024), using both the mapped RNAseq reads and the OrthoDB V11 Eukaryote protein database (Kuznetsov et al. 2023).

### Genome analyses

Genome completeness based on the predicted proteins was assessed with BUSCO 5.5.0 and the “Eukaryota_odb10” set of reference proteins. Genome statistics were generated using the GAG script version 2.0.1 (Hall et al. 2014). Orthofinder version 2.5.2 (Emms and Kelly 2015) was used to identify groups of orthologous genes between *D. pulchra* and detect red algal reference proteins (Table 1), enabling the detection of expanded gene families as well as lineage-specific genes.

**Table 1.**
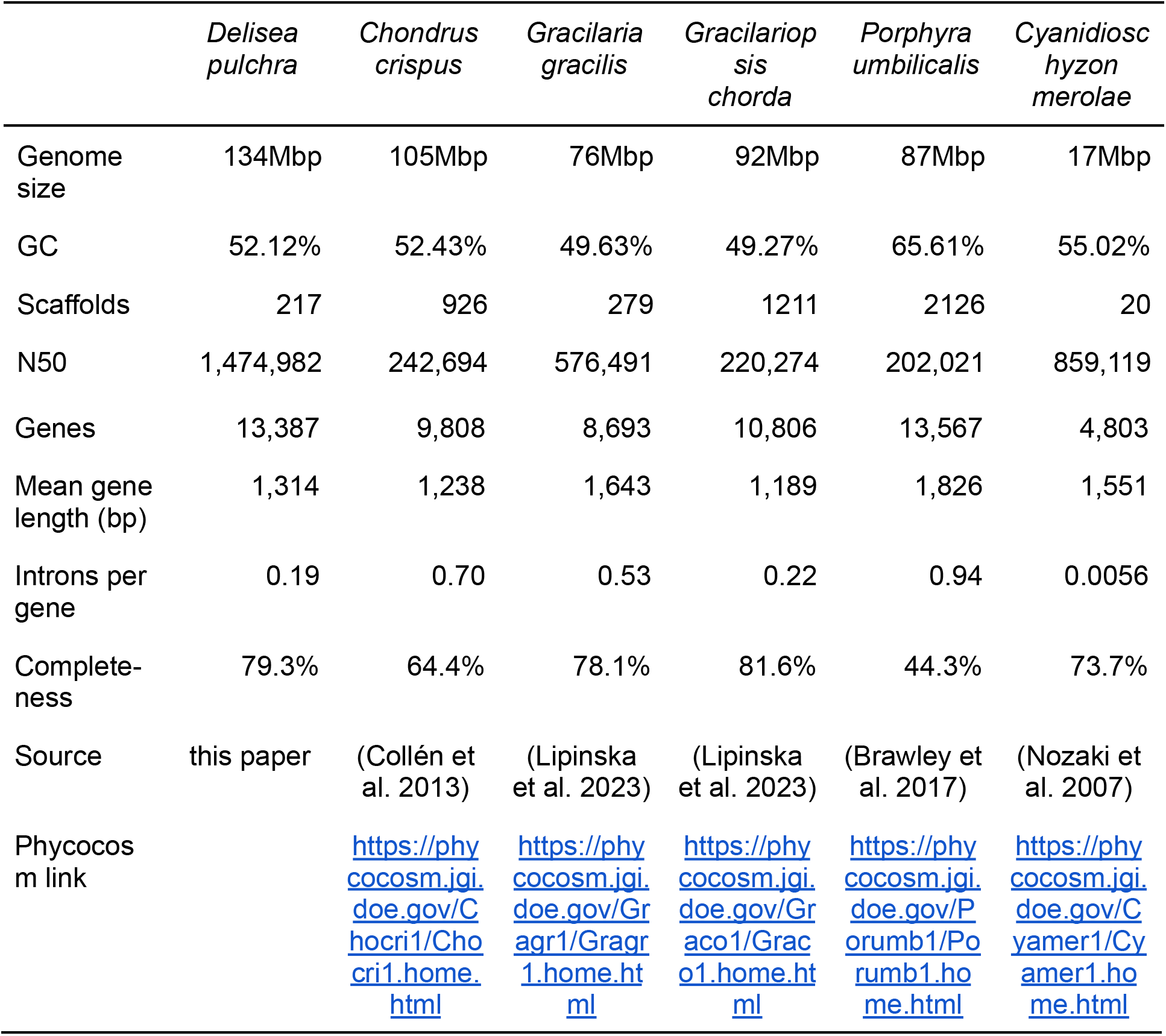
Overview of genome statistics of the *Delisea pulchra* and other selected red algal genomes. Completeness was calculated with BUSCO on the predicted proteins.

To annotate the proteome of *D. pulchra*, we utilised the ORSON pipeline (https://gitlab.ifremer.fr/bioinfo/workflows/orson) with a suite of state-of-the-art tools as well as with antiSMASH v8.0.4. Proteome completeness was evaluated by BUSCO 5.4.3 against the Eukaryota_odb10 lineage dataset (Manni et al. 2021). Functional annotations were generated with InterProScan 5.59-91.0 (Jones et al. 2014) and eggNOG-mapper 2.1.9 (Huerta-Cepas et al. 2019, Cantalapiedra et al. 2021). Subcellular targeting was predicted using HECTAR 1.3 (Gschlössl et al. 2008), and sequence similarities to the UniRef90 database (March 2022 release) were identified with Diamond 2.0.15 (Buchfink et al. 2014). The assembled organellar genomes were annotated via the CHLOROBOX web service on Jan 4th, 2024 (https://chlorobox.mpimp-golm.mpg.de/index.html) (Tillich et al. 2017, Greiner et al. 2019).

## Data availability

The *D. pulchra* genome assembly was deposited at the European Nucleotide Archive under ENA project accession PRJEB101077. It has furthermore been incorporated into the Rhodoexplorer portal (https://rhodoexplorer.sb-roscoff.fr/) at the Roscoff Biological Station (Lipinska et al. 2023). The *D. pulchra* genomic data can be explored and analyzed through a comprehensive web portal: https://abims-gga.sb-roscoff.fr/sp/delisea_pulchra. This user-friendly interface integrates several tools, including JBrowse (Buels et al. 2016) for visualizing the genome (with gene models, repeated elements, and RNA-seq data), GeneNoteBook (Holmer et al. 2019) for detailed functional annotation views, and ElasticSearch for quick gene lookups based on various criteria. BLAST-based sequence similarity searches against different *D. pulchra* sequence types are also available via SequenceServer (Priyam et al. 2019). Furthermore, the portal allows users to download all available *D. pulchra* datasets and provides links to external reference databases. The BEAURIS system (https://gitlab.com/beaur1s/beauris) was used to automate both the functional annotation process and the deployment of the entire *D. pulchra* genome portal.

## Results and Discussion

### Sequencing and Genome Assembly

The sequencing runs yielded a total of 29.15 million read pairs for the MiSeq run, corresponding to 4.37 Gbp of sequence data, and 1.03 million reads for the Sequel II run (average length 7.5 kb), corresponding to 7.76 Gbp of sequence data. The size of the assembled and cleaned *D. pulchra* genome was 134 Mbp, distributed among 271 contiguous sequences, with an N50 of 1.47 Mbp, and 13,387 protein-coding genes. 67.29% of the genome was composed of repeated elements, mainly retroelements, unclassified transposons, and DNA transposons (see Supplementary Table S1 for details). No ambiguous bases or gaps were found in the final nuclear genome assembly. The plastidial and the mitochondrial genomes were assembled into single, circular contigs with respective sequence lengths of 181.5 and 28.5 kb (illustrated in Supplementary Figure S1). Overall, genome size, assembly quality, and BUSCO completeness are on par with other long-read sequenced red algal model species and in the range of what is expected for red algae (see Table 1).

### Gene Structure

Studying the gene structure, it was found that the genes of *D. pulchra* have very few introns, only 0.19 introns per gene, a very low value for a multicellular eukaryote and lower than the other multicellular red algal species shown in Table 1, despite having a larger genome size. A low number of introns per gene has been found in all red algae hitherto studied (Borg et al. 2023) and is thought to be related to an ancestral genome reduction with loss of genes and introns in the rhodophytes (Qiu et al. 2015). *D. pulchra* is also a relatively gene-rich red alga with over 13,000 genes (Table 1).

### Orthofinder analyses

Orthofinder analyses (Supplementary Table S2) were performed to compare the predicted proteins of the *D. pulchra* nuclear genome with the selected reference genomes presented in Table 1. In total, 9,634 orthogroups (OGs) were detected, each containing between two and 201 proteins (see Supplementary Table S2). Of the 25 orthogroups that expanded most clearly in *D. pulchra, i*.*e*. those for which at least 20 more proteins were identified in *D. pulchra* than for any of the other examined red algae, 13 (52%) are completely unknown, 5 (20%) correspond to likely transposable elements that had not been detected by RepeatMasker, and 7 (28%) can be associated with putative functions (Supplementary Table S2). Prominent among these are OGs related to DNA methylation: Notably, OG0000000 comprises 193 copies in *D. pulchra* vs. a maximum of 6 in the other red algal genomes. This OG comprises probable methyltransferases of unknown specificity. Similarly, OG0000009 comprises probable DNA Methylases with 45 proteins in *D. pulchra* and a maximum of one in other red algae.

Another interesting group (although not within the top 25 orthogroups) is OG 30, which comprises Glutathione S-transferase-like proteins. *D. pulchra* has 15 while other red algae have 1-7. Similarly, we identified an OG of superoxide dismutases (OG0000041) found in over 20 copies in *D. pulchra* vs. a maximum of 2 in the other examined genomes. Lastly, some gene families involved in carbohydrate metabolism have also expanded in *D. pulchra*, and these genes are discussed in detail below.

### Extracellular matrix-specific genes and functions

The extracellular matrix (ECM) of *D. pulchra* remains undescribed in the literature, and genes related to the synthesis of ECMs were therefore examined manually. Other red macroalgae in the Bonnemaisoniales order, *Asparagopsis armata* (Haslin et al. 2000, Garon-Lardiere, S. 2004) and *Asparagopsis taxiformis* (Rodríguez Sánchez et al. 2023), have a mixture of complex sulfated galactans described as resembling agar and carrageenan in their ECMs. The genome of *D. pulchra* has several multi-copy gene families coding for carbohydrate-active enzymes of interest (Drula et al. 2022) predicted to be involved in the biosynthesis of agars and carrageenans, as described for the agarophyte red alga *Porphyra umbilicalis* and the carrageenophyte red alga *Chondrus crispus*. Glycosyl transferases (GTs) form the glycosidic linkages during polysaccharide biosynthesis, and *D. pulchra* has 4 GT7s, 2 GT8-GT64 double module enzymes, 19 GT14, and 3 GT47 members, which may be involved in sulfated galactan biosynthesis (Collén et al. 2013, Ficko-Blean et al. 2015, Brawley et al. 2017, Lipinska et al. 2020). We also identified 22 GT2 members, 3 of which are predicted to encode for cellulose synthase (CESA). Genes encoding for predicted sulfotransferase (ST) enzymes were identified as 2 ST1, 9 ST2, 1 ST3, and 2 Gal3OST. Since the ST2 and Gal3OST families are carbohydrate-specific, these are good candidates for agar or carrageenan sulfation. Unique to red algae are the galactose-6-sulfurylases, responsible for the formation of the 3,6-anhydro-bridge in 3,6-anhydro-galactose, a sugar found only in agars and carrageenans and responsible for their gelling properties (Genicot-Joncour et al. 2009, Lipinska et al. 2020,2020, Rhein-Knudsen and Meyer 2021, Chevenier et al. 2023). One partial hit on a galactose-2,6-sulfurylase type I was found, and 2 putative galactose-2,6-sulfurylases type II were identified in *D. pulchra*. Three GH16 enzymes were found; one red algal GH16 enzyme from *C. crispus* has been characterised as a porphyranase (Manat et al. 2022), but other enzyme functions in the GH16 family include κ-carrageenase (Potin et al. 1991) and agarase activities (Jam et al. 2005). More unexpectedly, a sulfatase with ∼28 % sequence identity to the characterised bacterial S1_12 choline-sulfatase was found (Van Loo et al. 2018, Stam et al. 2023). This enzyme type is not generally found in red algae, but the model is supported by RNAseq reads, well integrated into a scaffold containing other algal gene models, and exhibits >30% identity with bacterial genes. It might be involved in small-molecule desulfation or matrix remodeling.

### Defense genes

While *D. pulchra* is well known for its capacity to produce halogenated furanones (Dworjanyn et al. 2006) and for the impact these compounds have on interactions with its microbiome (Harder et al. 2012), very little is known about the biochemical pathways that lead to their production. One lead comes from a paper by Sandy et al. (2011), who demonstrated that algal-type vanadium bromoperoxidases (vBPOs) can act on 4-pentynoic acid to form a bromofuranone. Another study by Thapa et al. (2020) showed that *A. taxiformis* possesses a gene cluster including three vBPOs and an NAD(P)H oxidase (NOX), implicated in bromoform production.

In the genome of *D. pulchra*, we identified three homologs of the algal-type vBPOs characterized in *A. taxiformis* (Thapa et al. 2020), four bacterial-type vanadium haloperoxidases (vHPOs; homologs to NapH enzymes producing chlorinated antibiotics; McKinnie et al. 2018), and eight NOX genes (Supplementary Table S4). We also identified 19 heme-dependent haloperoxidases (hHPOs), which have distinct enzymatic mechanisms and evolutionary histories compared to vBPOs (Hofrichter and Ullrich 2006). Lastly, we identified a gene cluster comprising one NOX, two of the putative vBPOs, and four heme-dependent bromoperoxidases (hBPOs) (Figure 2A). Any of these genes is potentially involved in furanone synthesis in *D. pulchra*, but none of them is specific to this species or even to the Bonnemaisoniales.

**Figure 2.**
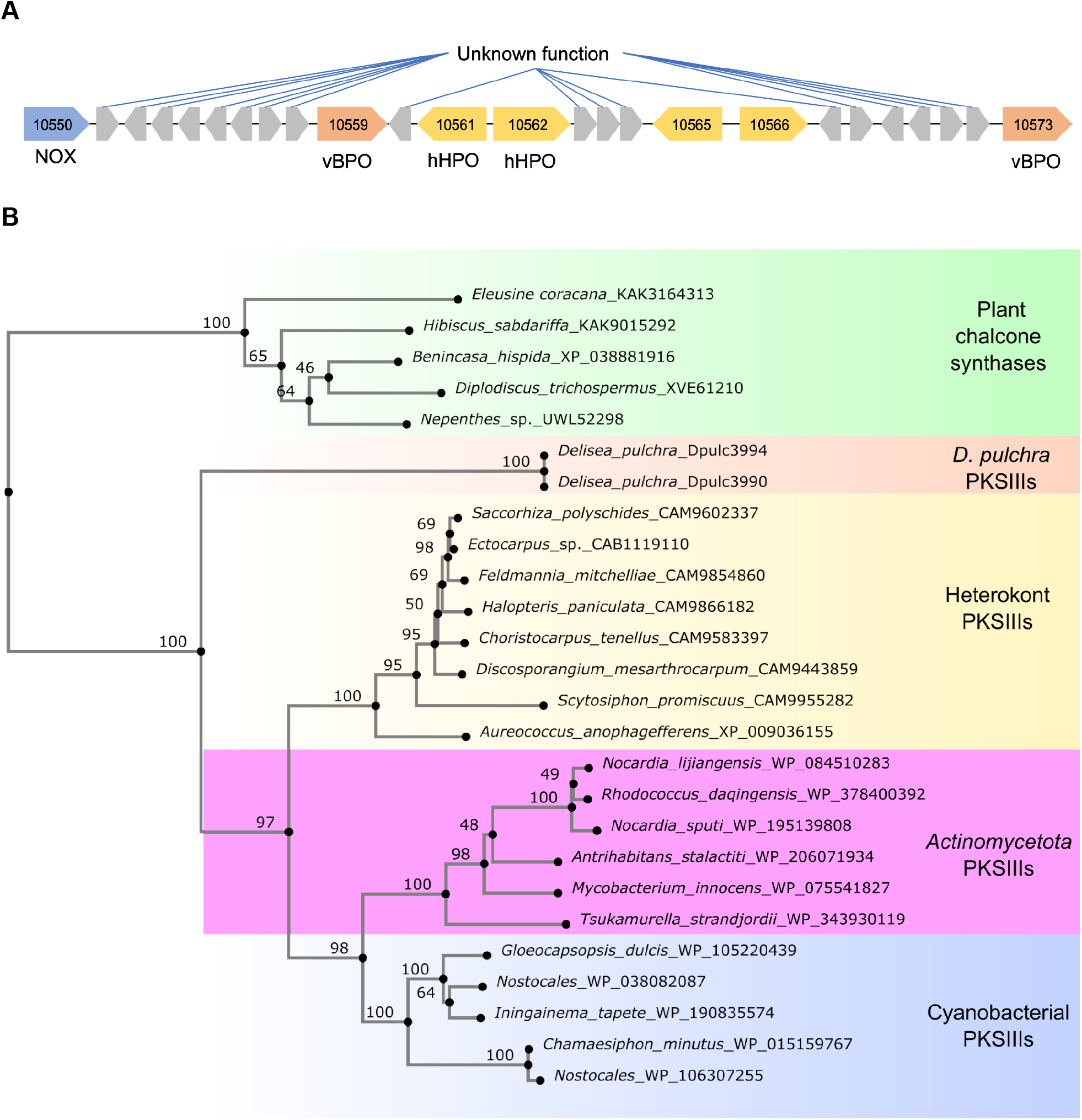
A) A representation of the gene cluster that is potentially involved in bromoform production. The genes are not to scale. Genes are indicated by their numeric identifiers; full locus tags follow the format DpulcXXXXX. B) The phylogenetic tree is based on *D. pulchra* PKSIII proteins, as well as selected BLAST hits from the RefSeq select database. The sequences were aligned using MAFFT v7 and the G-INS-i algorithm. A neighbour-joining tree was then calculated using the WAG substitution model and 100 bootstrap replicates on all 287 gap-free sites of the alignment. The tree was visualised using phylo.io.

One hypothesis for the formation of furanones specifically in *D. pulchra* and related algae is the availability of distinct substrates for these enzymes to act on. In this context, we identified two 99.5% identical, adjacent type III polyketide synthase (PKS) genes that are unique to *D. pulchra* and phylogenetically distant from both bacterial and other eukaryotic PKSs (Figure 2B). While unsaturated fatty acids analogous to 4-pentynoic acid remain the classical candidate precursors of furanones, the cyclicized lactons produced by PKSs also consitute plausibe substrastes and warrant further investigation.

In addition to halogenated furanones, we identified eight terpene biosynthetic gene clusters (Supplementary Table S4) in the antiSMASH analysis of the *D. pulchra* genome, suggesting that terpenes are also part of its defense repertoire, as has been described in *Laurencia pacifica* and some other Florideophyceae (Carter-Franklin et al. 2003).

## Conclusion

Here we present a high-quality genome assembly for the red macroalga *Delisea pulchra*, which is a valuable addition to the limited genomic resources currently available for Rhodophyta. This highly contiguous, complete, and functionally annotated genome provides a solid basis for molecular and functional research on this species.

The genome reveals several unique features, including expanded gene families associated with DNA methylation, oxidative stress responses, and carbohydrate metabolism. These features may be related to complex extracellular matrix biosynthesis and to the alga’s capacity to respond to environmental challenges, including host–pathogen interactions. Identification of an unusual choline sulfatase highlights the potential for discovering new metabolic adaptations in this species. Similarly, identification of *D. pulchra*-specific type III polyketide synthases and several haloperoxidases provides promising candidates for future studies of furanone biosynthesis. These findings, combined with the presence of multiple terpene biosynthetic clusters, highlight the alga’s metabolic investment in its unique chemical defenses.

The *D. pulchra* genome will facilitate targeted research into the genetic basis of chemical defence, disease susceptibility, and microbial interactions, as well as comparative analyses across red algal lineages. By integrating this resource into the Rhodoexplorer platform, we aim to make this genome more accessible and facilitate comparative studies across red algal lineages.

## Acknowledgments

The work presented in this paper was funded by the CNRS/INSB via the “International call 2022 - Marine Biology” project DIPLOMAT. We thank the Roscoff Bioinformatics platform, ABIMS (http://abims.sb-roscoff.fr), and the Genomer platform, respectively part of the Institut Français de Bioinformatique (ANR-11-INBS-0013), of EMBRC-France (financially supported by the Investments of the Future program, ANR-10-INSB-02), and of the Biogenouest network, for their support. We are grateful to the Gentyane platform at Clermont-Ferrand for library preparation and PacBio sequencing.

## Supplementary Material

**Supplementary Table S1.** Repeated elements detected with RepeatMasker 2 in the *Delisea pulchra* genome.

|  | number of elements | length occupied (np) | percentage of sequence (%) |
| --- | --- | --- | --- |
| **Retroelements** | 16935 | 34976966 | 26.09 |
| SINEs: | 0 | 0 | 0.00 |
| Penelope | 0 | 0 | 0.00 |
| LINEs: | 98 | 91161 | 0.07 |
| CRE/SLACS | 0 | 0 | 0.00 |
| L2/CR1/Rex | 31 | 49834 | 0.04 |
| R1/LOA/Jockey | 0 | 0 | 0.00 |
| R2/R4/NeSL | 0 | 0 | 0.00 |
| RTE/Bov-B | 0 | 0 | 0.00 |
| L1/CIN4 | 0 | 0 | 0.00 |
| LTR elements: | 16837 | 34885805 | 26.02 |
| BEL/Pao | 0 | 0 | 0.00 |
| Ty1/Copia | 4204 | 5389046 | 4.02 |
| Gypsy/DIRS1 | 12633 | 29496759 | 22.00 |
| Retroviral | 0 | 0 | 0.00 |
| **DNA transposons** | 24018 | 23168952 | 17.28 |
| hobo-Activator | 73 | 40496 | 0.03 |
| Tc1-IS630-Pogo | 1349 | 847089 | 0.63 |
| En-Spm | 0 | 0 | 0.00 |
| MuDR-IS905 | 0 | 0 | 0.00 |
| PiggyBac | 92 | 103016 | 0.08 |
| Tourist/Harbinger | 1046 | 713432 | 0.53 |
| Other | 0 | 0 | 0.00 |
| **Rolling-circles** | 282 | 267187 | 0.20 |
| **Unclassified:** | 43098 | 32075325 | 23.92 |
| **Total interspersed repeats:** |  | 90221243 | 67.29 |
| **Small RNA:** | 154 | 122439 | 0.09 |
| **Satellites:** | 0 | 0 | 0.00 |
| **Simple repeats:** | 313 | 23044 | 0.02 |
| **Low complexity:** | 0 | 0 | 0.00 |

**Supplementary Table S2.** Gene counts per orthogroup as identified by Orthofinder. The second tab contains the gene identifiers for each species and Orthogroup.

**Supplementary Table S3.**
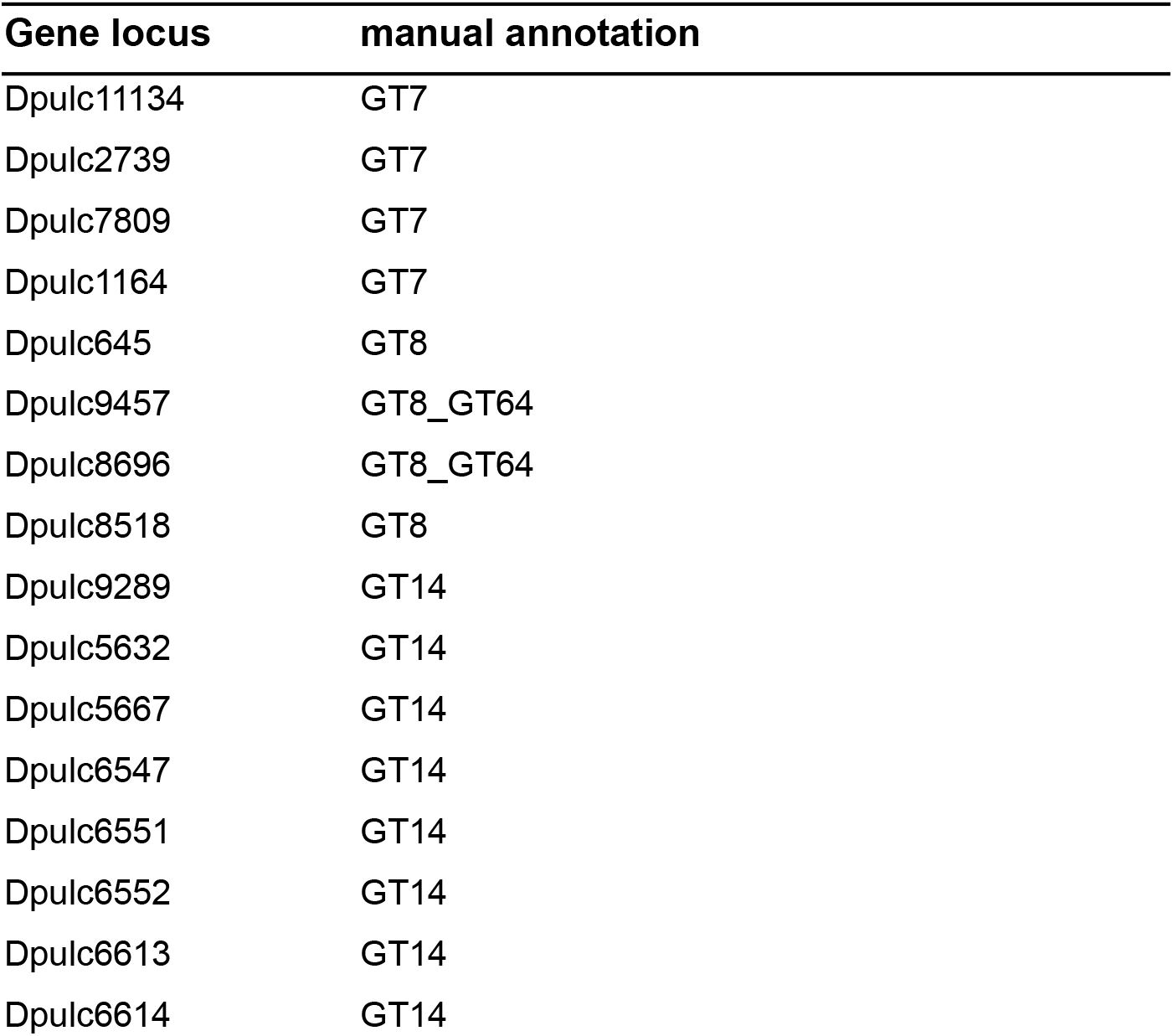
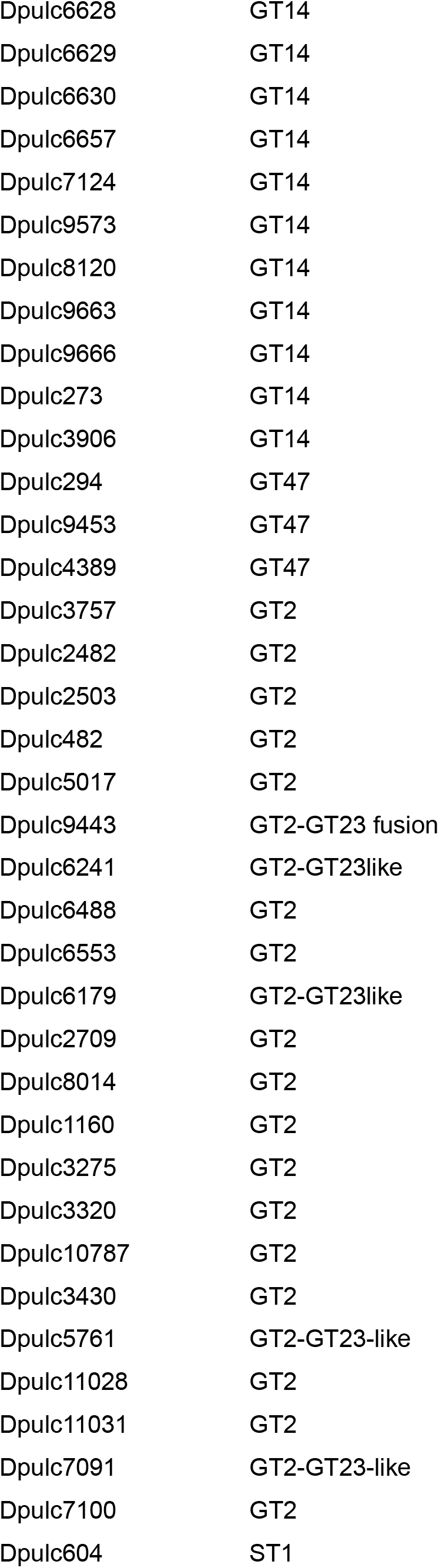
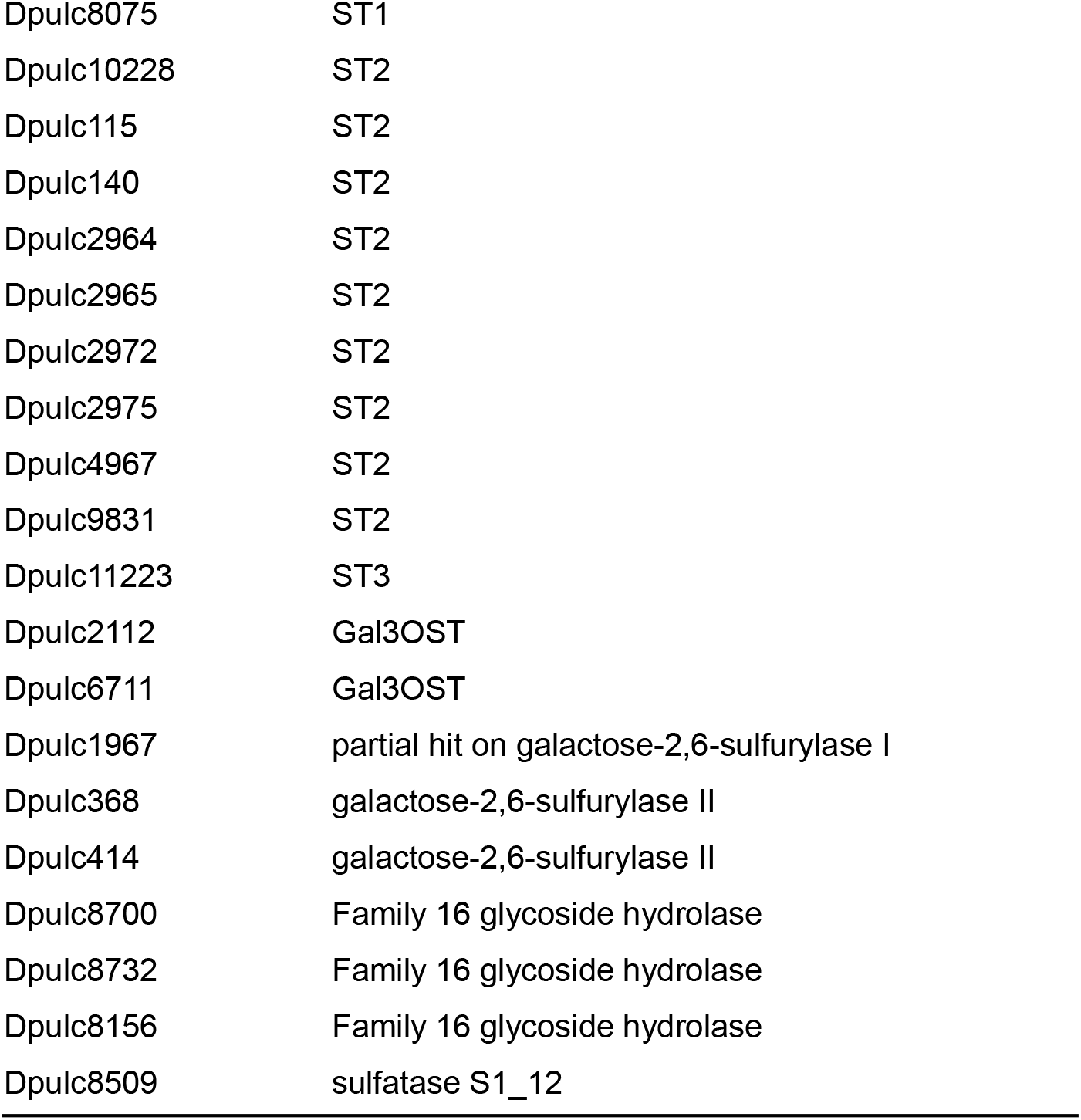
Manual annotation of proteins involved in carbohydrate metabolism and extracellular matrix biosynthesis in *Delisea pluchra*.

**Supplementary Table S4.** Manual annotation of proteins involved in defense in *Delisea pluchra*.

| Gene accession | manual annotation |
| --- | --- |
| Dpulc6207 | Vanadium haloperoxidase ("algal-type") |
| Dpulc10559 | Vanadium haloperoxidase ("algal-type") |
| Dpulc10573 | Vanadium haloperoxidase ("algal-type") |
| Dpulc8818 | Vanadium haloperoxidase ("bacterial-type") |
| Dpulc8819 | Vanadium haloperoxidase ("bacterial-type") |
| Dpulc9238 | Vanadium haloperoxidase ("bacterial-type") |
| Dpulc11260 | Vanadium haloperoxidase ("bacterial-type") |
| Dpulc4028 | Heme Peroxidase |
| Dpulc4092 | Heme Peroxidase |
| Dpulc4094 | Heme Peroxidase |
| Dpulc4100 | Heme Peroxidase |
| Dpulc5312 | Heme Peroxidase |
| Dpulc5315 | Heme Peroxidase |
| Dpulc5318 | Heme Peroxidase |
| Dpulc5782 | Heme Peroxidase |
| Dpulc6562 | Heme Peroxidase |
| Dpulc6565 | Heme Peroxidase |
| Dpulc6951 | Heme Peroxidase |
| Dpulc6952 | Heme Peroxidase |
| Dpulc8914 | Heme Peroxidase |
| Dpulc8917 | Heme Peroxidase |
| Dpulc10561 | Heme Peroxidase |
| Dpulc10562 | Heme Peroxidase |
| Dpulc10565 | Heme Peroxidase |
| Dpulc10566 | Heme Peroxidase |
| Dpulc11574 | Heme Peroxidase |
| Dpulc5785 | Haloalkane dehalogenase |
| Dpulc5786 | Haloalkane dehalogenase |
| Dpulc5792 | Haloalkane dehalogenase |
| Dpulc5793 | Haloalkane dehalogenase |
| Dpulc5794 | Haloalkane dehalogenase |
| Dpulc5940 | Haloalkane dehalogenase |
| Dpulc8411 | NAD(P)H oxidase (NOX) |
| Dpulc6202 | NAD(P)H oxidase (NOX) |
| Dpulc10550 | NAD(P)H oxidase (NOX) |
| Dpulc4130 | NAD(P)H oxidase (NOX) |
| Dpulc8084 | NAD(P)H oxidase (NOX) |
| Dpulc1434 | NAD(P)H oxidase (NOX) |
| Dpulc4129 | NAD(P)H oxidase (NOX) |
| Dpulc3081 | NAD(P)H oxidase (NOX) |
| Dpulc1302 | NAD(P)H oxidase (NOX) |
| Dpulc3990 | Type III Polyketide synthase (PKS III) |
| Dpulc3994 | Type III Polyketide synthase (PKS III) |
| Dpulc3767 | 3-ketoacyl-CoA synthase (KAS) |
| Dpulc4516-Dpulc4521 | Terpene biosynthetic cluster 1 |
| Dpulc11236-Dpulc112352 | Terpene biosynthetic cluster 2 |
| Dpulc8399-Dpulc8407 | Terpene biosynthetic cluster 3 |
| Dpulc5558-Dpulc5565 | Terpene biosynthetic cluster 4 |
| Dpulc6072-Dpulc6081 | Terpene biosynthetic cluster 5 |
| Dpulc1454-Dpulc1465 | Terpene biosynthetic cluster 6 |
| Dpulc5357-Dpulc5363 | Terpene biosynthetic cluster 7 |

**Supplementary Figure S1.**
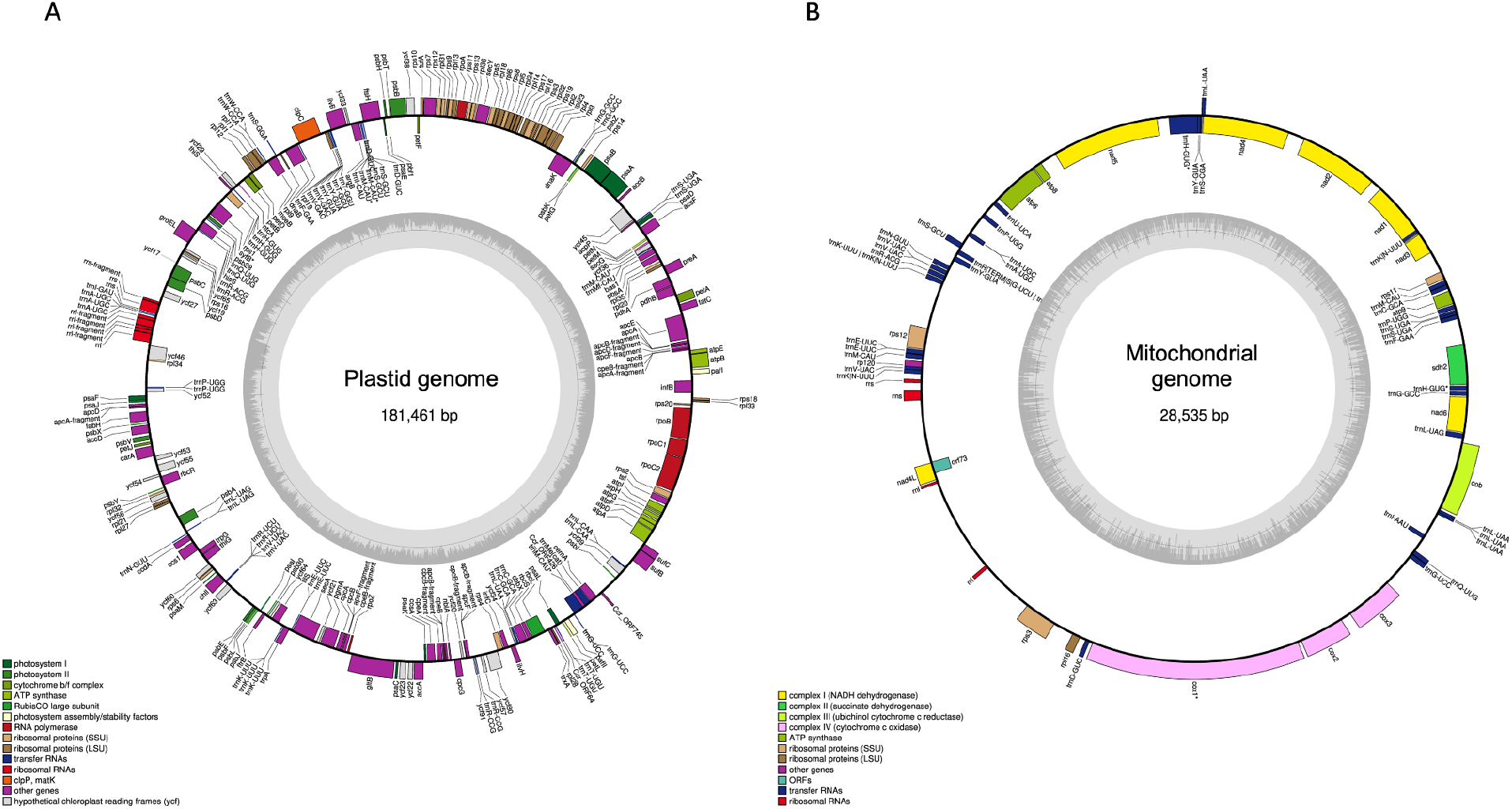
Visualisations of the plastidial (A) and the mitochondrial (B) genomes of *Delisea pulchra*.

